# SciGeneX: Enhancing transcriptional analysis through gene module detection in single-cell and spatial transcriptomics data

**DOI:** 10.1101/2024.03.19.585667

**Authors:** Julie Bavais, Jessica Chevallier, Lionel Spinelli, Serge A. van de Pavert, Denis Puthier

## Abstract

The standard pipeline to analyze scRNA-seq or spatial transcriptomics data focuses on a gene-centric approach, which overlooks the collective behavior of genes. However, cell populations should be viewed as intricate combinations of activated and repressed pathways. Thus, a broader view of gene behavior would provide more accurate information on cellular heterogeneity in single-cell or spatial transcriptomics data. Here, we described SciGeneX, a R package implementing a neighborhood analysis and a graph partitioning method to generate co-expression gene modules. These gene modules, which can be shared or restricted between cell populations, collectively reflect cell populations, and their combinations are able to highlight specific cell populations, even rare ones. SciGeneX was also able to uncover rare and novel cell populations which were not observed before in spatial transcriptomics data of human thymus. We show that SciGeneX outperforms existing methods on both artificial and experimental datasets. Overall, SciGeneX will aid in unraveling cellular and molecular diversity in single-cell and spatial transcriptomics studies. The R package is available at https://github.com/dputhier/scigenex.

## Introduction

Recent advances in high-throughput next-generation sequencing (NGS) technologies have revolutionized the transcriptomics field, offering detailed information on individual cells [1]. Among these technologies, single-cell RNA-seq (scRNA-seq) has emerged as a powerful and widely used method for investigating cellular heterogeneity within complex tissues [2,3]. Even more recently, spatial transcriptomics (ST) has emerged as a groundbreaking technology, allowing for the mapping of gene expression in tissues. ST is anticipated to revolutionize biology by providing unparalleled insights into the intricate interplay of genes and cells within their native microenvironments [4,5]. Over 1000 tools have been developed to analyze scRNA-seq [6] such as *Scanpy* [7], *DUBStepR* [8], *singleCellHaystack* [9], or *Hotspot* [10]. Seurat [11] is still the predominant choice in current studies [12] and has become a standard R library for scRNA-seq and ST data analysis [13]. The standard pipeline to analyze these types of data typically begins with pre-processing steps aimed at removing non-biological sources of variation. First, quality control is performed to identify and remove low quality cells and ambient RNA [14]. Subsequently, cell counts are normalized to mitigate technical noise or biases while preserving biological variation [15]. A feature selection step is then used to extract informative genes [16] which are subsequently reduced to a latent space using principal component analysis (PCA). Unsupervised cell clustering, usually based on Euclidean distance [17], is then performed using the latent space. Afterwards, differential gene expression analysis is conducted to identify genes that are expressed in specific clusters [18], enabling groups of cells to be assigned to distinct cell types or states. Cells are subsequently visualized in a two dimensional space using t-distributed stochastic neighbor embedding (t-SNE) [19] or uniform manifold approximation and projection (UMAP) [19]. A similar pipeline is used to cluster spots in ST data.

In contrast to gene clustering, which can be evaluated using functional enrichment, the cell or spot clustering approach used in the classical pipeline does not rely on any biological indicator. Identifying the optimal cell partitioning requires multiple empirical clustering attempts, along with the identification of differentially expressed genes (DEGs). Moreover, DEG quality is intricately linked to the cell clustering quality. For instance, the failure to identify a rare cell population during cell clustering prevents marker gene identification for this population [20]. Additionally, the dual use of gene information to generate cell or spot clusters and to identify DEGs leads to double-dipping, inducing a high type I error rate [21]. Finally, the standard approach to scRNA-seq and ST analysis places its primary emphasis on dissecting cellular or spot intricacies through an examination of genes on an individual basis. However, this gene-centric analysis, while informative, falls short of encapsulating the whole comprehensive complexity inherent in cellular biology. The standard pipeline, by focusing on a gene-by-gene study, overlooks the collective behavior of genes and may inadvertently neglect those that, while less variable individually, could collectively define critical cellular states. This limitation becomes particularly evident when dealing with genes expressed in rare populations, which often elude detection using the conventional methodology, resulting in the inadvertent exclusion of entire cell populations from analysis.

In contrast, cell populations are the outcomes of intricate differentiation and activation processes shaping epigenetic landscapes that selectively activate or repress specific genomic loci. Viewing cell types and states as intricate combinations of activated and repressed pathways offers a more holistic perspective. A promising alternative to the standard pipeline is to first cluster genes into co-expression modules and then group the cells according to their modules usage [22]. It is within this biological context that we developed SciGeneX (Single-cell informative Gene eXplorer), an R package implementing an algorithm to generate co-expression gene modules. Unlike the standard pipeline, SciGeneX does not limit itself to a gene-centric analysis but instead delves into the broader panorama of gene co-expression. It opens up new possibilities for understanding cellular dynamics and heterogeneity. This paradigm shift reflects a deeper commitment to capturing the intricate relationships between genes, ultimately leading to more meaningful insights into the biological underpinnings of scRNA-seq and ST data.

The SciGeneX algorithm is an adaptation of the DBF-MCL algorithm which we previously developed in the context of microarray analysis [23]. SciGeneX performs feature selection using a neighborhood analysis to select co-expressed genes across cell populations. It then partitions these into gene modules using the Markov Cluster Algorithm [24]. We benchmarked the SciGeneX gene module approach against classical methods using a set of artificial and real datasets. We found that SciGeneX substantially outperformed conventional methods such as DISP and VST, from the Seurat R package, to retrieve informative genes. Using both a small (3K peripheral blood mononuclear cells) [25] and a large dataset (∼70K thymocytes) [26] containing both mature and differentiating cells, we show that the gene modules generated by SciGeneX are highly resolutive and enriched in specific biological processes. These modules, reminiscent of specific cell markers, cell states and numerous signaling pathways can be combined to reveal biologically relevant cell groups independently of the classical scRNA-seq analysis pipeline. Interestingly, the relevance of our procedure is strengthened by the ability of SciGeneX to reveal previously undetected rare cell populations and associated marker genes. Finally, we demonstrate that the SciGeneX approach translates to Visium spatial transcriptomics (ST) data and can elucidate biologically meaningful topological molecular profiles. To store the co-expression gene modules, we have implemented an S4 R class named ‘ClusterSet’. This implementation empowers the users to access various functionalities to visualize, characterize, and manipulate the discovered gene modules.

Overall, our results indicate that SciGeneX introduces a paradigm shift to scRNA-seq and ST data analysis. Through its innovative co-expression module generation, SciGeneX outperforms conventional methods, demonstrating superior efficacy in identifying and extracting informative genes. It provides highly resolved gene modules, enriched in specific biological processes, capable of revealing previously undetected rare cell populations and associated markers. SciGeneX’s adaptability to spatial transcriptomics data further underscores its significance in advancing comprehensive biological insights.

## Results

### Overview of the SciGeneX pipeline

The SciGeneX algorithm operates through a sequence of four key steps: (i) elimination of genes that do not co-express with other genes, (ii) construction of a gene’s neighborhood graph, (iii) identification of modules of co-expressed genes, (iv) filtering of modules of co-expressed genes.

In the initial phase, it employs a density-based-filtering (DBF) approach, removing genes that do not exhibit significant co-expression with their neighbors across cells. This is achieved by comparing the gene-gene distance matrix, often based on the Pearson correlation coefficient, to a null hypothesis of random gene distribution. The distances to the K^th^ nearest neighbors (DKNN) are computed, and a threshold is determined to distinguish co-regulated genes (see Methods). In the second step, the algorithm constructs a gene’s neighborhood graph using two available methods: “closest_neighborhood” and “reciprocal_neighborhood.” The former establishes edges based on the size (S) of the closest neighborhood of each gene, while the latter considers reciprocal relationships within a neighborhood of size K. Finally, the algorithm employs the Markov Cluster Algorithm (MCL) to partition co-expressed genes into modules. MCL simulates a random walk through the graph, identifying densely connected regions that represent distinct biological programs. A subsequent filtering step is applied to handle singleton clusters and ensure the reliability of the co-expression modules generated by the algorithm.

### The SciGeneX algorithm outperforms classical feature selection methods

To determine whether the DKNN used by SciGeneX accurately ranks genes according to the information they carry, we applied SciGeneX on 100 artificial datasets (Fig. 2A). We used SPARSim [27] to generate artificial datasets containing noise and a predetermined number of DEGs distributed into 4 gene modules. We used a range of maximum fold-change values to simulate DEG under various conditions. As shown in Supplementary Data, Figure S1, above a maximum fold change of 10, these artificial gene modules result in increasingly segregated cell populations. Ten independent artificial datasets were generated for each maximum fold-change value, resulting in a total of 100 datasets. Finally, we compared the ranking score of SciGeneX with six existing methods: DISP (dispersion) [28], VST (variance stabilizing transformation) [29] and sctransform [30] from the Seurat package [11] as well as M3Drop [31], singleCellHaystack [9] and BigSur [32].

**Figure 1.**
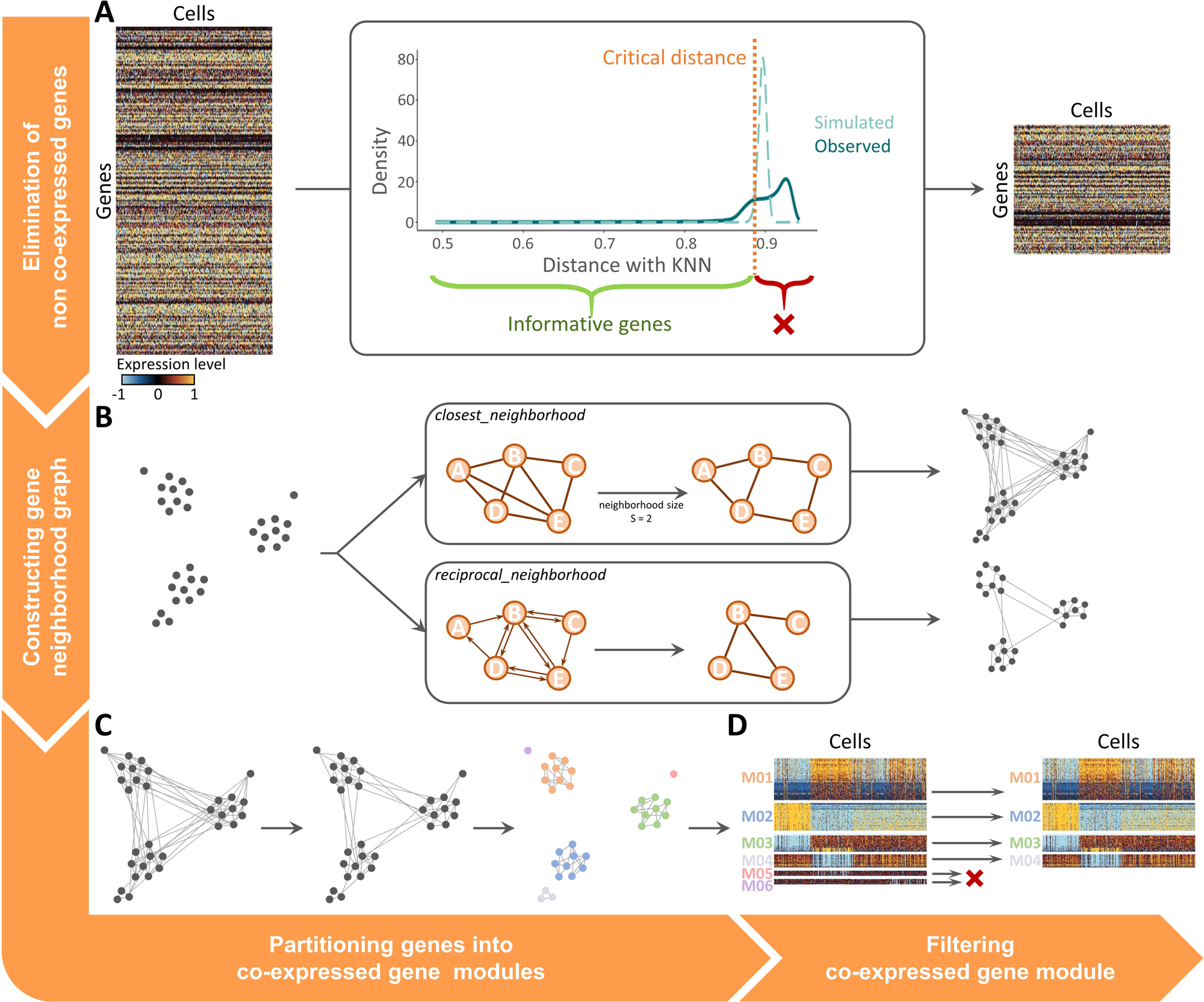
Overview of SciGeneX algorithm. The SciGeneX algorithm consists of four consecutive steps. (**A**) The algorithm first eliminates non co-expressed genes employing a density-based filtering approach. It involves comparing the gene-gene distance matrix, typically using the Pearson correlation coefficient, against a null hypothesis of random gene distribution. Distances with the K nearest neighbor (DKNN) are computed, and a threshold is established to eliminate genes that do not co-express with other genes. (**B**) Then, the SciGeneX algorithm constructs a gene’s neighborhood graph. Two methods are available. The *closest_neighborhood* method establishes edges based on the size (S) of each gene’s closest neighborhood while the *reciprocal_method* considers reciprocal relationships within a neighborhood of size K. (**C**) Gene selected are then clustered into co-expression modules using the MCL algorithm. (**D**) Finally, co-expression modules are filtered typically based on module size or standard deviation.

**Figure 2.**
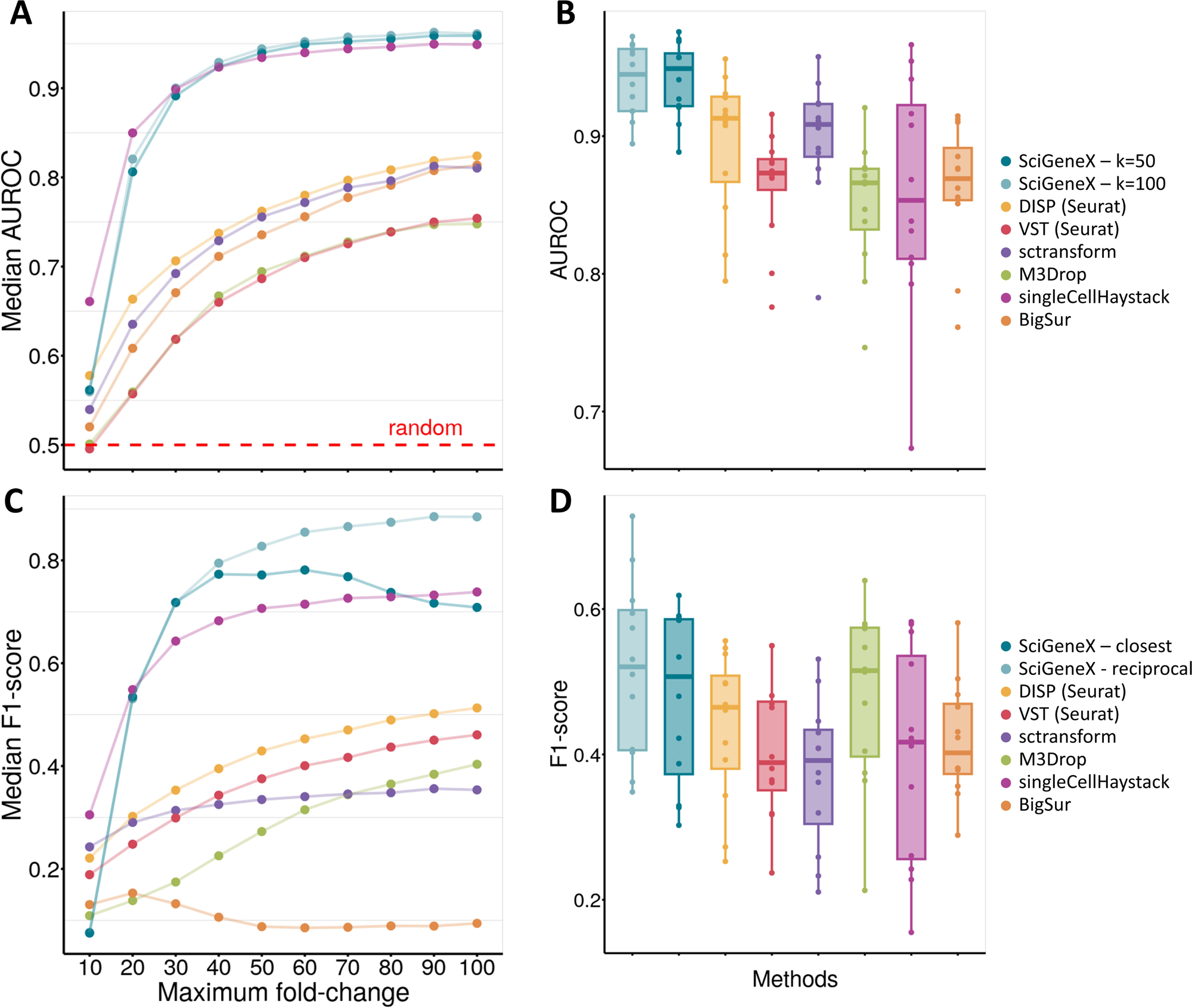
Performance evaluation for finding true DEGs in artificial and experimental datasets. Performances were evaluated through two widely used scores, AUROC and F1-score using both artificial and experimental datasets. AUROC was used to evaluate the performance of the neighborhood analysis of SciGeneX for two K values, K=50 (dark blue) and K=100 (light blue) while F1-score was used to evaluate the ability of both SciGeneX classifiers, *closest_method* (dark blue) and *reciprocal_method* (light blue). These results were compared with six methods, DISP (yellow) and VST (red) from the Seurat R package, sctransform (purple), M3Drop (green), singleCellHaystack (pink) and BigSur (orange) (**A**, **C**) Scatter plots displaying median AUROC (**A**) and median F1-score (**C**) computed across artificial datasets. The medians were based on 10 replicated over a maximum fold-change range (10 to 100, in increments of 10). (**B**, **D**) Scatter plots showing AUROC (**B**) and F1-score (**D**) computed across of set of experimental datasets from the Tabula Muris consortium generated on 12 tissues.

Using the area under the receiver operating characteristic (AUROC) curve, we measured the ability of the SciGeneX neighborhood analysis (neighborhood size, K, set to 50 and 100) and the six other methods to rank DEGs. The neighborhood analysis on artificial datasets displayed higher AUROC medians compared to all the other methods throughout almost all artificial datasets (Fig. 2A). Only singleCellHaystack showed higher median AUROC values while close to the ones of SciGeneX in artificial datasets generated with low maximum fold-change (10 to 30). Of note, for a maximum fold-change of 10, genes do not properly segregate cell populations (Supplementary Data, Figure S1) and poor performances are thus expected.

While artificial datasets are particularly useful to set up a method, they may not adequately mimic the counts from experimental datasets. In contrast, experimental datasets lack the predetermined knowledge about which genes are truly informative (*i.e.* that differs from noise). To overcome this limitation, we used the well characterized Tabula Muris [3] datasets and considered the cell population markers obtained through DEGs identification as the true informative genes out of the whole gene set (Fig. 2B). As the DEG list may differ according to the approach, we used four popular methods including *Wilcoxon* (Fig. 2B,D), *bimod* [33], *MAST* [34], and *t.test* (Supplementary Data, Figure S2). DEGs were selected with a minimum log2 fold-change of 0.5 (*i.e* linear fold change of ∼1.4) and a minimum adjusted p-value of 0.05. Similar as for the AUROC scoring in the artificial datasets, SciGeneX yielded higher AUROC values for experimental datasets when compared to the six other methods (Fig. 2B). This indicated that the SciGeneX neighborhood analysis better distinguished DEGs from noise compared to the other methods. Overall, the DKNN used in the SciGeneX neighborhood analysis outperforms the compared methods in ranking informative genes within artificial and experimental datasets.

After evaluating the DKNN, we computed the performances of the algorithm as a whole: gene filtering, gene partitioning and module filtering. To do this, we computed the F1-score that integrated both precision (*i.e.* the proportion of genes predicted as DEGs that are actually DEGs) and recall (*i.e.* the proportion of actual DEGs that were predicted as DEGs). As SciGeneX implements two methods, *closest_neighborhood* and *reciprocal_neighborhood*, to create the graph subsequently partitioned by the MCL algorithm, the gene list produced by these two methods were assessed using F1-scores. Performances were then compared with the previously mentioned methods. When run on artificial datasets, both SciGeneX algorithms displayed higher F1-score medians in general except with a maximum fold change of 10 (Figure 2C). Only singleCellHaystack showed higher F1-score while remaining close to SciGeneX results when the maximum fold-change is low (between 10 to 20). The performance of the *reciprocal_neighborhood* method tended to drop as the maximum fold-change increased, however, only singleCellHaystack showed higher scores. When tested on 12 experimental Tabula Muris datasets, both SciGeneX algorithms showed higher median F1-score, with the exception of the M3Drop method which showed higher median F1-score (0.515) while remaining close that of the *closest_neighborhood* SciGeneX method (0.507) (Fig. 2D). Same results were observed with the three other approaches used to identify the DEGs (Supplementary Data, Figure S2).

Our results indicate that both SciGeneX algorithms are powerful methods for distinguishing informative genes from non-informative genes in both artificial and experimental datasets. They both showed consistent high AUROC and F1-score across all modalities making it an attractive tool to select informative genes.

### Combination of co-expressed gene modules reveals cell populations

One of the key advantages of the SciGeneX algorithm is its ability to partition informative genes into co-expression modules using the MCL algorithm. To evaluate the ability of the MCL algorithm to produce gene modules from selected informative genes which would overlap with specific biological functions, we used the PBMC3k [25] dataset as a gold standard. This dataset, available on the 10x Genomics website, contains 2638 cells and 13714 genes. When employing the standard Seurat tutorial, cells were classified into 9 reference cell populations using Louvain classification. The SciGeneX neighborhood analysis identified 3340 genes while the MCL algorithm partitioning generated 377 co-expression modules. These gene modules were filtered based on the number of genes and the number of cells (n=2) expressing at least 60% of the genes from the corresponding modules. After filtering, a set of 977 informative genes clustered into 16 co-expression modules were generated. Following gene partitioning into modules, cell ordering was performed using the hierarchical clustering algorithm based on genes from co-expression modules generated by the SciGeneX algorithm. This classification successfully highlighted the previously identified cell populations (Figure 3A). Interestingly, as shown in Figure 3A, gene modules were reminiscent of previously known cell types. Module M04 was found to be specifically active in a previously characterized platelet population, while module M16 demonstrated specific expression in dendritic cells (Fig 3a). Additionally, several modules demonstrated shared expression among multiple cell populations. For instance, module M05 exhibited high expression in natural killer cells and moderate expression in CD8+ T lymphocytes. Module M06 displayed a high level of expression in both populations. On the other hand, module M07 showed predominant expression in T lymphocyte populations, including CD8+ and CD4+ T lymphocytes (Figure 3A).

**Figure 3.**
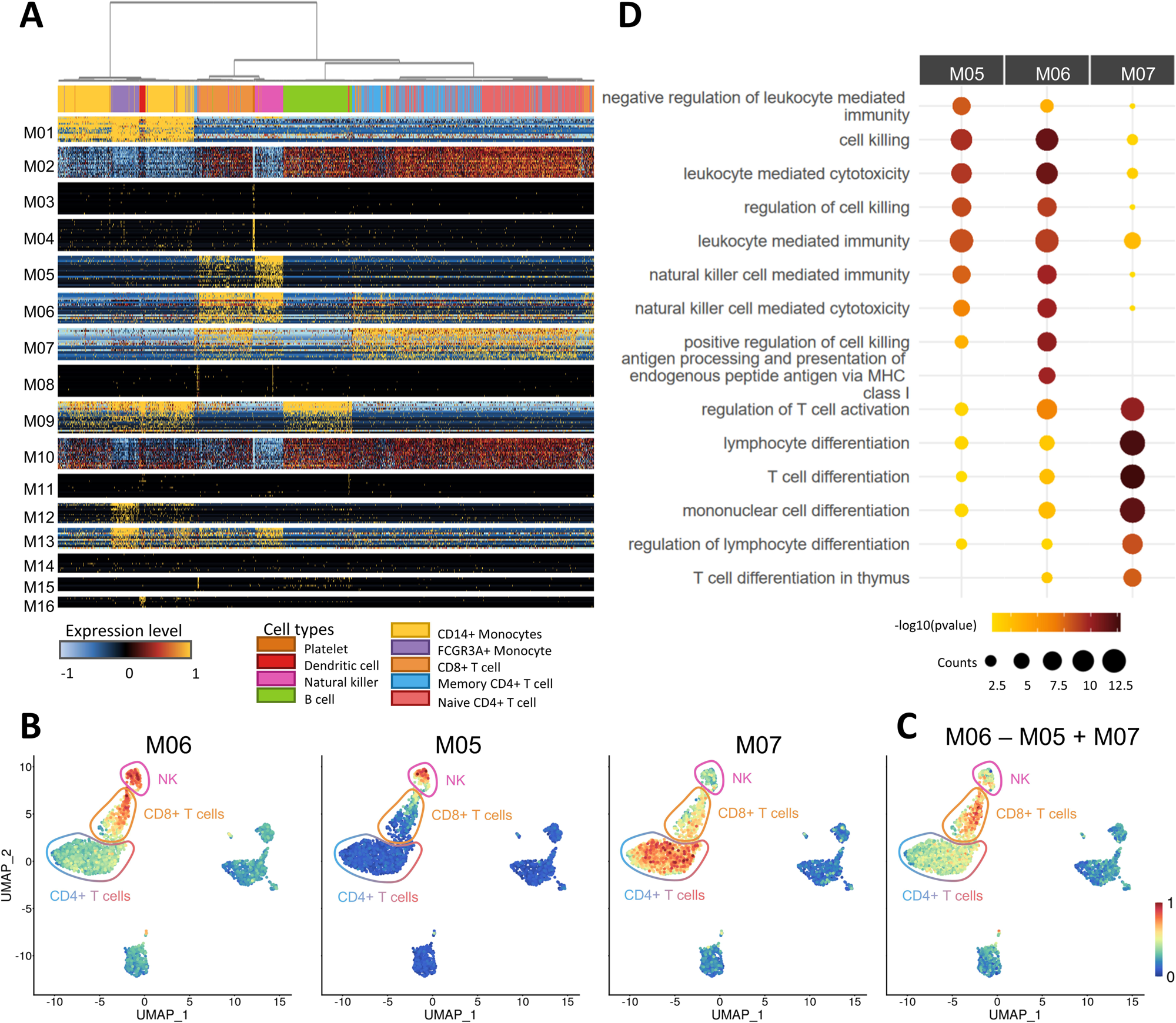
Combinations of gene modules generated by the SciGeneX algorithm reveal cell populations in relation to biological processes. (**A**) Heatmap displaying normalized expression levels of the top 20 genes within each co-expression module generated by the SciGeneX algorithm. Cells have been ordered using a hierarchical clustering algorithm. The labels of the cell populations, as defined in the original Seurat results, are shown on the top of the heatmap. (**B**) UMAP plot of PBMCs colored by their AUCell score for module M06, M05 and M07 from left to right. (**C**) The AUCell scores of module M06 were subtracted from those of module M05 and added to those of module M07 to obtain a composite score, which is plotted on the UMAP. (**D**) Dotplot showing the Gene Ontology (GO) enrichment analysis for gene module M05, M06 and M07. The top 5 enriched GO terms are displayed for each gene module, providing insights into the biological processes associated with the previously mentioned co-expression modules.

Cell populations are the result of differentiation processes that lead to setting up of epigenetic landscapes that direct the activation or repression of specific loci. In this regard, cell types and states can be viewed as combinatorics of activated and repressed pathways. We hypothesized that these repressed/activated pathways could be mimicked by combinations of gene module activities that would reveal underlying cell types. To summarize module activity, we computed AUCell scores for each of them across all cells and projected them on a UMAP. We can visualize individual module activities (*i.e.* AUCell scores) (Figure 3B) and sum up the activities of shared modules M06 and M07 subtracted by the activity of shared module M05 (Figure 3C). This can be interpreted as looking for cells with high levels of module M06 and M07 while excluding cells with high levels of module M05. Interestingly, such a combination revealed a new module with high specificity in CD8+ T lymphocytes which was supported by functional enrichment analysis (Figure 3D). Indeed, module M07 exhibits enrichment in biological processes related to T cell differentiation, a function shared between CD4 and CD8 T cells. Module M06 was found to be enriched in genes related to cell killing and cytotoxicity, which are functions shared between CD8 T cell and NK. While the functional enrichment of module M05 closely resembled that of module M06, a more detailed examination of the gene list revealed a significant presence of NK-specific markers including various killer cell immunoglobulin like receptor (such as *IR2DL3* and *KIR3DL2*) and killer cell lectin like (including *KLRC1*, *KLRD1*, and *KLRF1*).

Altogether, these results demonstrate the ability of the SciGeneX algorithm to capture co-expressed gene modules associated with both specific and shared expression between cell populations which are almost all significantly associated with specific terms from biological processes (Supplementary Data, Figure S3). We show that these modules can be used alone or combined to reflect cell populations and assist downstream cell clustering analysis.

### Co-expression modules reveal rare cell populations

While the original analysis on the PBMC3k dataset revealed 9 cell populations, the gene modules revealed by SciGeneX suggested a more intricate situation. Close inspection of the hierarchical clustering in Figure 3A revealed that the expression of modules M11 and M15 were restricted to two different and yet small cell groups. Applying a two-class k-means clustering to the expression data of module M11 and M15 elucidated these populations (Figure 4A). Interestingly, cells expressing genes from module M11 clustered together in the UMAP, and this was also true for cells expressing genes from module M15. To further characterize these populations, we interrogated the ImmuNexUT database that provides Count Per Million (CPM) values for genes across a diverse set of FACS-sorted immune cell populations. Genes from module M11 were predominantly expressed in plasmacytoid dendritic cells (Figure 4C). This observation was in line with the fact that *LILRA4*, *CLEC4C*, *SERPINF1*, and *ILR3A* have been previously reported as markers of this population in several studies referenced in the PanglaoDB database [35]. Similarly, genes from module M15, including *TNFRSF17*, *UGT2B17*, and *MZB1,* were mainly active in plasmablasts which is consistent with the fact that they are known plasmablasts markers in the PanglaoDB database.

**Figure 4.**
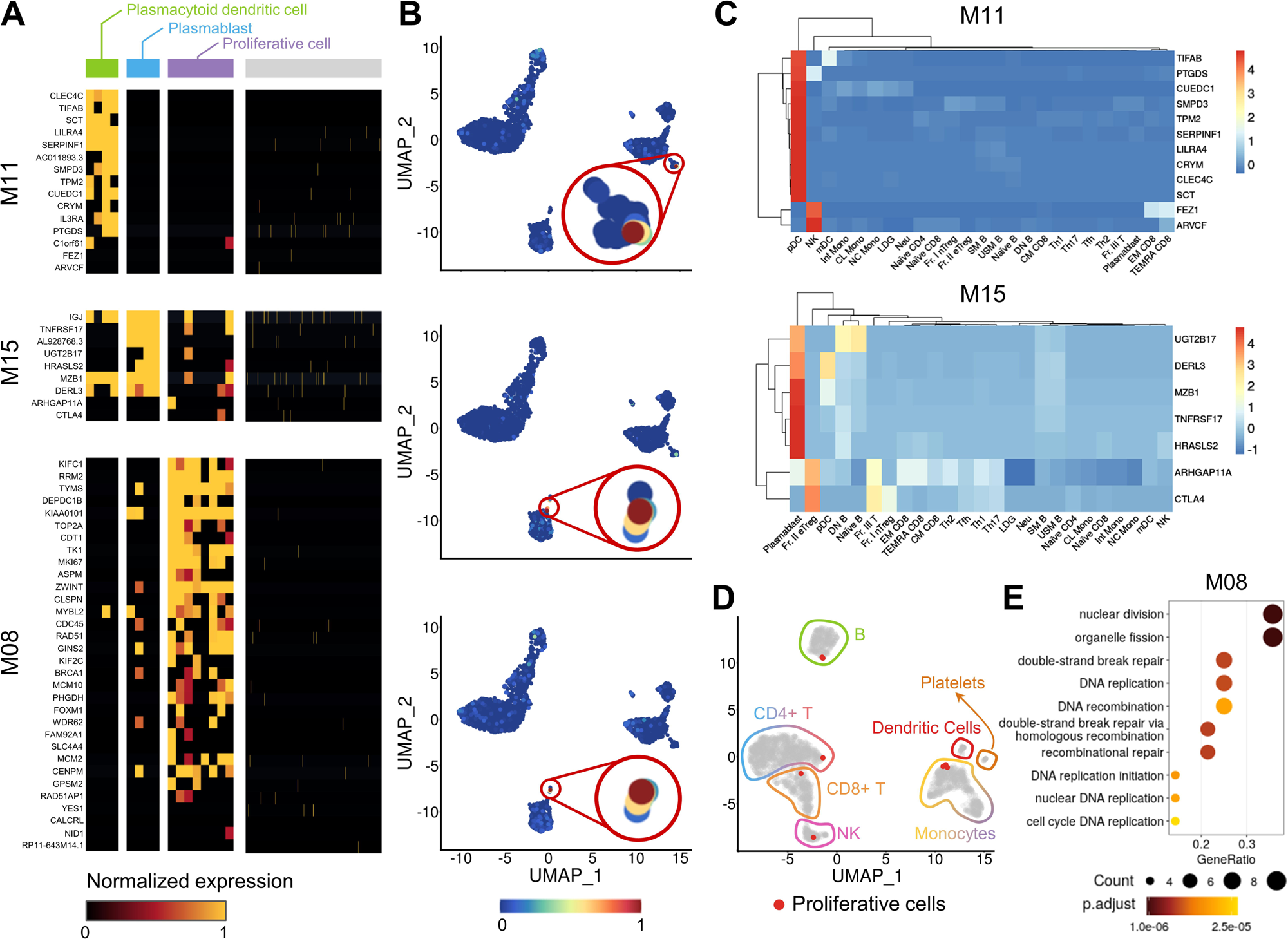
Identification of rare cell populations using co-expression modules. (**A**) Heatmap representing normalized expression levels within co-expression modules M08, M11 and M15 generated by the SciGeneX algorithm. Cells have been partitioned using K-Means clustering method. The cell clusters are indicated at the top of the heatmap with 4 cells in green, 4 cells in blue, 8 cells in purple, and 2622 in gray. (**B**) UMAP plot of PBMCs colored by their AUCell score for co-expression modules M08, M11 and M15 (top to bottom). (**C**) Heatmap displaying the CPM values for immune cell types from ImmuNexUT, focusing on genes from co-expression modules M11 and M15. (**D**) UMAP representation of PBMCs with gene expression regressed for genes belonging to co-expression module M08. Proliferative cells are highlighted in red. (**E**) Dotplot showing the results of GO enrichment analysis for gene module 8. The top 5 enriched GO terms are displayed providing insights into the biological processes.

Moreover, using a hierarchical clustering analysis, we identified genes from module M08 (Figure 4A) that were expressed in distantly related populations (Figure 3A). A two-class k-means clustering was also used to reveal these populations. Surprisingly, these cells appeared to be co-localized within a B-cell population in the UMAP (Figure 4B). Gene ontology analysis unveiled that genes from module M08 were related to cell proliferation (Figure 4E). We thus regressed expression data with genes from module M08 and performed a new gene reduction and UMAP representation. Contrary to what was initially thought, these cells did not cluster exclusively within the B cells population but distributed among several populations. This highlighted very tiny sub-populations of proliferating natural killer cells, CD14+ monocytes, and CD4+ T lymphocytes (Figure 4D).

Our results showcase the ability of the SciGeneX algorithm to reveal extremely rare yet functionally relevant cell populations or cell states using a dataset that has already been extensively explored.

### Co-expression modules reveal cellular heterogeneity in a continuum of differentiating T cells

The PBMC3k dataset previously analyzed was characterized by a low complexity and the presence of terminally differentiated cells. We directed our focus toward a substantially larger dataset comprising developing cells. To this end, we used the “Human T cell trajectory” dataset [26] containing 76,994 cells (∼30 times more than the PBMC3k dataset) that were classified into 16 main cell populations, offering a continuous spectrum of differentiating cells. Using the *reciprocal_neighborhood* SciGeneX method, we identified a total of 2821 informative genes, distributed across 71 co-expression modules, with an average of 40 genes within each module (ranging from 197 to 8) (See Supp_file_Gene_Modules_Thymus_Single_Cell.xls and Supplementary Data, Figure S4). In this context, the *reciprocal_neighborhood* method was preferred as it was found to produce more balanced gene modules (ranging from 197 to 8 genes) compared to the *closest_neighborhood* method (data not shown).

The modules generated by the SciGeneX algorithm reflected the diverse stages characterizing thymic T-cell development (Figure 5). Among them, module M11 is characterized by the presence of key factors associated with very early T-cell development, including the pivotal transcription factor NOTCH1 and its target HES1 as well as the transcription factor RUNX2 [36]. Module M03 is associated to the double-positive (DP) stage with prominent markers such as CD8A, CD8B, CD4, CD52, CCR9, and the transcription factors LEF1, TCF7, TCF12, SATB1, in addition to the VDJ recombination actors RAG1 and RAG2 [37]. Module M06, in turn, contains more mature T-cell markers, including CD5, CD6, CCR7, CD28, accompanied by interferon-inducible genes such as IFI44L, IFITM1, IFITM2, HLA-A, HLA-B, HLA-C, HLA-F, and IRF7.

**Figure 5.**
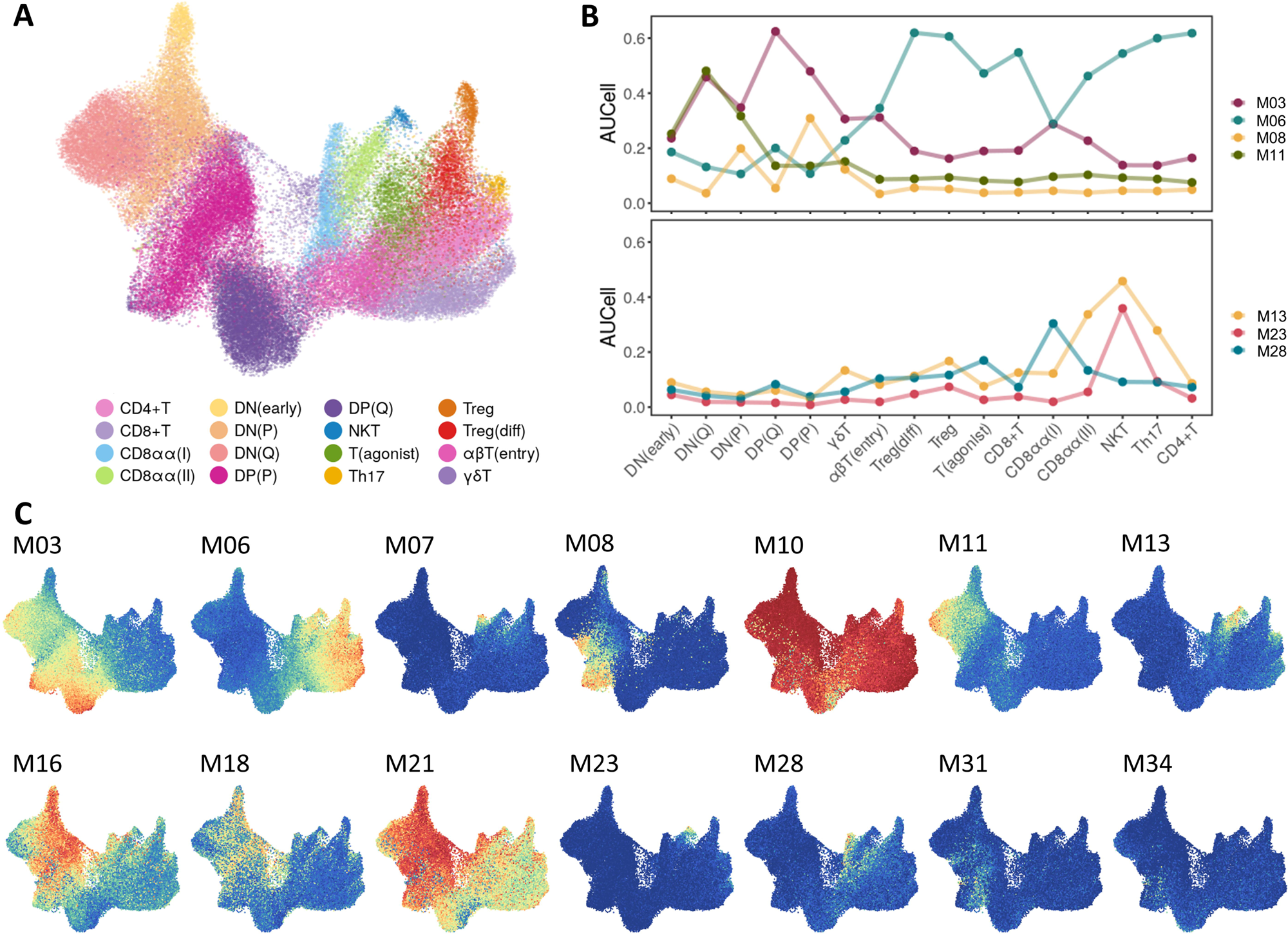
Co-expression modules reveal differentiation processes in a continuum of developing T cells. (**A**) UMAP plot of T cell differentiation stages from human thymus. Cells are colored by the cell types defined in the original study. (**B**) Evolution of the expression of gene modules across the differentiation of T cells. The top graph shows modules M03, M06, M08 and M11 related to cellular functions and the bottom graph shows the modules M13, M23 and M28 related to known differentiation stages. (**C**) UMAP plot colored by AUCell scores for co-expression modules found by the SciGeneX algorithm.

T-cell differentiation is marked by the regulation of several biological processes. These events are mirrored by various gene modules: module M08 that corresponded to a proliferative state (*i.e.* MKI67, TOP2A, BIRC5, CCNB1, CCNB2, CCNF,…), module M10 reflected protein synthesis (*i.e.* RPL10, RPL11, RPL12, RPL13, RPL14, RPL15,…) while modules M16, M18, and M21 reflected the developmentally regulated usage of various genes related to oxidative phosphorylation (*i.e.* COX17, COX5A, and COX8A, NDUFA1B, NDUFB7, NDUFS6…) (Figure 5C). Moreover, T-cell differentiation is marked by profound chromatin rearrangements. These events are particularly highlighted by module M31 containing 20 genes related to histones and localized on chromosome 6 (*i.e.* HIST1HA, HIST1HB, HIST1HD, HIST1HE, HIST1H2AG,…) or module M34 composed also of 18 genes coding for histones that harbor a more restricted expression program (Figure 5C).

The set of 71 modules generated by SciGeneX represents a comprehensive valuable source of information. For instance, we noted that within the dataset described by JE Park et al., three distinct CD8aa+T cell populations, GNG4+CD8aa+T(I) cells, ZNF683+CD8aa+T(II) cells, and an EOMES+ CD8aa+NKT-like population, were recently identified. SciGeneX successfully recovered modules closely reminiscent of these three populations, each characterized by highly specific genes encompassing surface markers, chemokines, chemokine receptors, and transcription factors, providing valuable insights for a more refined population characterization. For example, module M28, specific to GNG4+CD8aa+T(I) cells, was comprised of 30 marker genes, including 3 known markers, GNG4, CREB3L3, PDCD1, along with 27 additional genes such as NFATC1, FZD3, CD200, and CD82. Interestingly, module M07 was even more specific to the GNG4+CD8aa+T(I) cell population, with a slightly distinct profile. It contained the chemokine XCL1, known to recruit XCR1+ dendritic cells, as recently described in the analysis of JE Park *et al.* Our analysis also brought to light the presence of the XCL2 chemokine in this module, alongside several additional markers, including CD72 and the microRNA MIR3142HG. Module M13 was synonymous to the ZNF683+CD8aa+T(II) cell population characterized by a mixed αβ and γδ lineage as previously reported [26]. This module, comprising 77 genes, revealed several markers associated to the γδ T-cell receptors (e.g TRDV1, TRDV2, TRG-AS1, TRGC2, TRGV10, TRGV2, TRGV7) alongside the marker CD84 and various killer cell lectin-like and killer cell immunoglobulin-like receptors (e.g KIR3DL2, KLRB1, KLRC3) and the RUNX3 transcription factor. Finally, module M23, comprising 38 genes, was found to be specific to the EOMES+ CD8aa+NKT-like population. Notably, this population exhibited specific surface markers (CD160, FCGR3A, IL18RAP, KLRC1, KLRC4, KLRD1, FASLG) and demonstrated the production of several cytokines, including CCL3, CCL3L3, CCL4L2, and IFNG.

Altogether, our results highlight the ability of the SciGeneX algorithm to create highly biologically meaningful gene modules that depict the complex processes underlying T-cell differentiation.

### Scigenex applied to spatial transcriptomics data allow in depth resolution of tissue molecular architecture

The advent of spatial transcriptomics technologies has ushered in a new era in biological research, paving the way for unprecedented insights into the spatial organization of gene expression in tissues, while simultaneously presenting formidable computational challenges to harness the full potential of this revolutionary approach [38,39]. In this context, some solutions have been recently developed to reveal spatially regulated gene patterns [9,10,40]. In order to assess the ability of the SciGeneX algorithm in detecting co-expression modules in spatial transcriptomics data, we ran the SciGeneX algorithm on a recently published Visium dataset interrogating the transcriptome of a human thymus section (sample GSM6281326 from GEO Serie GSE207205) [41]. In this study, the authors previously stained thymic slices using H&E that is known for its ability to delineate the two primary regions within the thymus: the cortex situated in the organ’s outer layer and hosting early T-cell precursors, and the medulla which contains more mature thymocytes (Figure 6A). The Seurat reference analysis identified 7 spot clusters that are depicted in Figure 6B and corresponding DEG (not shown).

**Figure 6.**
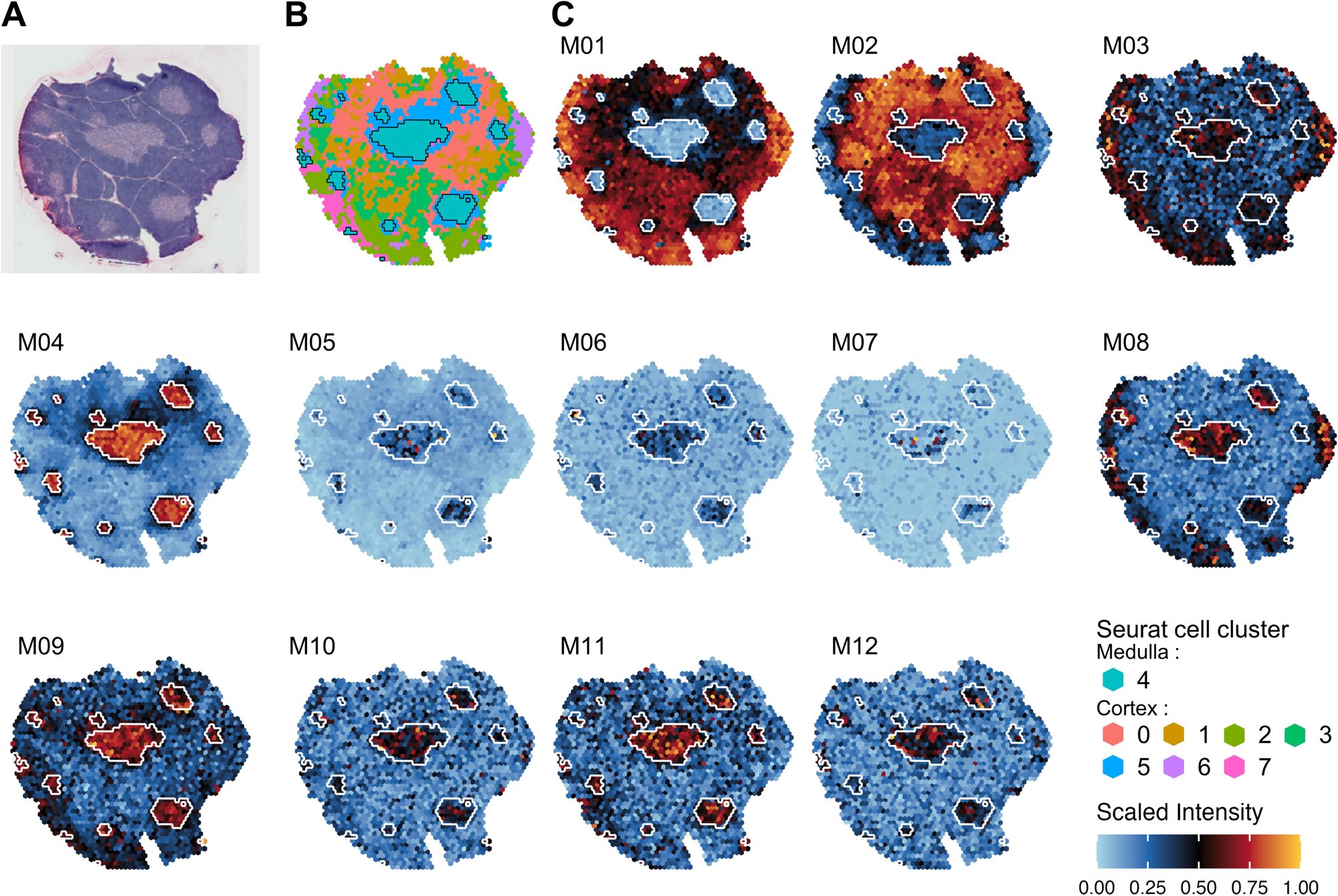
Co-expression modules generated by the SciGeneX algorithm on spatial transcriptomics of a human thymus section. (**A**) H&E image of human thymus and matching spots annotated into 8 topological areas. Medullary areas appear as internal zones with less intense labeling. (**B**) Spot clusters as identified by the *FindClusters()* Seurat method (default parameters). The regions surrounded by a thin black line corresponds to the Seurat spot cluster associated with the medulla (Seurat spot cluster #4 in legend). (**C**) Scaled intensity per spot for 12 co-expression modules generated by the SciGeneX algorithm. To ease comparison, the Seurat spot cluster #4 is also depicted with a thin white line in all panels from (**C**).

Using Scigenex a set of 21 gene modules (containing at least 8 genes) corresponding to a total of 1402 genes (mean number of genes per module: 70.1) were revealed (Supplementary Data, Figure S5 and Supp_file_Gene_Modules_Thymus_Spatial_Transcriptomics.xls). The spatial distribution of these co-expression modules were highly concordant with the known architecture of the thymus. As an example, modules M01 and M02 were found to be highly expressed in the outer cortex. Surprisingly, while the medulla was represented by a unique spot cluster in Seurat analysis (spot cluster #4), Scigenex was able to reveal a far more complex scenario. Indeed, numerous modules (Figure 6C and Supplementary Data, Figure S6) with spatially restricted patterns in the medulla of with expression shared between the cortex and medulla were found (e.g. M3 to M14, M16 to M21). Most of the gene modules could be unambiguously associated with functions known to be active in the thymus (Supplementary Data, Figure S7). This encompasses GO terms associated with T-cell development “V(D)J recombination and alpha-beta T-cell activation” (gene module M01), “positive regulation of lymphocyte activation” (gene module M04), “Chemokine activity” (gene module M06), “response to interferon−alpha” (gene module M11) or “ribosome assembly” (M15). Importantly, our method was sensitive enough to detect several rare populations of highly specialized thymic epithelial cells (TEC) including: muscle mimetic medullary TEC (gene module M07, enriched for “striated muscle contraction”) known to specifically express skeletal muscle genes (MYOG, TNNT1-3, MYL4…), corneoTEC (gene module M05, enriched for “epidermal cell differentiation”) known to express keratinocytes or corneocytes markers (*e.g.* DMKN, LY6D and SPINK5) or AIRE^+^ mTECs (gene module M06, enriched for “CCR chemokine receptor binding”) [42]. Altogether, these results demonstrate that, applied to Visium datasets, SciGeneX algorithm can provide a highly resolutive overview of the cellular diversity. Our results also indicate that even with a low resolution technology (spot diameter 50µm), SciGeneX is able to identify spatially variable modules expressed in widespread or very rare mimetic cell populations [43].

## Discussion

In this study, we introduced SciGeneX which proposes a paradigm shift revolutionizing the scRNA-seq or ST analysis process by looking at the broader panorama of gene co-expression without prior cell clustering. Using gene neighborhood analysis and graph clustering algorithm, SciGeneX shows an innovative perspective by generating co-expression gene modules providing a comprehensive overview of the underlying biological processes within scRNA-seq or ST datasets. In contrast to conventional methods, SciGeneX aligns with the Waddington’s model [44], relying on the dynamic interplay of cellular states shaped by the activation or repression of biological pathways. Thus, cellular populations could be characterized by the activation and repression of various pathways, some of which may be specific or shared among different cell types.

In our study, SciGeneX has shown its ability to provide a very rich molecular overview of underlying cell populations. Indeed, it generated co-expression gene modules that collectively mirror well established cell populations. These gene modules can be either shared among multiple populations or selectively expressed in specific ones. For example, the expression of cytotoxic machinery components (such as perforins and granzymes in modules 5 and 6 of the PBMC3k dataset) is well-known to be highly confined to T cells and NK cells. However, while it serves as a recognized hallmark of cytotoxic cells, it is not exclusive to T cells or NK cells. Moreover, cellular populations could be characterized by the activation and repression of various pathways. Thus we showed that a combination of co-expression gene modules is able to highlight specific cell populations. For example, with a combination of 3 gene modules in the PBMC3k dataset, we were able to specifically highlight a population of CD8+ T cells. Conversely, certain modules can accurately detect the presence of exceptionally rare populations (e.g., plasmacytoid dendritic cells and plasmablasts) expressing highly specific markers. These populations with distinct markers were previously undetected using the Seurat pipeline, highlighting several limitations of conventional approaches.

Another benefit of Scigenex is its potential utility to provide biological cues to assist in accurately defining cell populations within a dataset. Indeed, while Scigenex does not offer a direct solution for cell clustering, it can contribute to decision-making by offering insights into activated or repressed pathways using heatmap visualization and mapping expression of specific or combined gene modules on a UMAP. This is crucial, as inaccurately defined cell populations can result in heterogeneous clusters, leading to reduced statistical power.

The SciGeneX algorithm also gave very promising and robust results when analyzing spatial transcriptomics data (Visium). Spatial transcriptomics is a rapidly evolving field with several concurrent but also complementary technologies. The SciGeneX algorithm was able to uncover very rare and recently discovered mimetic mTEC populations in ST dataset [45] which were not described in the original study [41]. As spatial transcriptomics continues to establish itself as a standard method, our algorithm could serve as a highly effective approach for investigating cellular diversity within spatial dimensions.

Additional datasets (3 Visium and several single-cell datasets) were also tested (data not shown). Across all the tested conditions, SciGeneX consistently uncovered highly relevant gene modules (note that a Visium brain dataset example is also provided in the accompanying vignette of the R package). Nevertheless, due to dataset diversity, it would be impractical to establish a one-size-fits-all set of optimal parameters. While this may seem like a limitation, users have the flexibility to adjust key parameters such as neighborhood size, the inflation parameter of the MCL algorithm, or filtering steps. Regarding filtering steps, we frequently achieved excellent results by simply choosing either the cluster size or standard deviation. However, the package also offers filtering options based on additional criteria, including silhouette similarity score, Dunn index, and connectivity score as proposed by the clValid R package [46]. Interestingly, SciGeneX is also intended to be a more general container of gene modules within the R ecosystem. In this way the ClusterSet object may be created from a Seurat object or from an expression matrix and gene list. This makes it fully compatible with other methods including *M3Drop* [31], *BigSur* [32], *DUBStepR* [8], *singleCellHaystack* [9], *Seurat or Hotspot* [10]. Once loaded, gene modules obtained from other methods may be manipulated easily by taking advantage of the ClusterSet methods.

In summary, SciGeneX is an approach of choice for unraveling cellular and molecular diversity in the fields of single-cell approaches and spatial transcriptomics. Its ability to offer a complete molecular snapshot of cell populations is particularly exciting for researchers keen to uncover hidden information in their data. The tool’s accessibility, adaptability and compatibility with other methodologies makes it a valuable asset in these research fields.

## Key Points

- We introduce SciGeneX, a R package to generate co-expression gene modules in single-cell RNA-sequencing and spatial transcriptomics data. It employs a gene neighborhood analysis coupled with a network clustering method.
- Different from the standard approach, SciGeneX focuses on the collective behavior of genes and provides initial insights into scRNA-seq and spatial transcriptomics data before cell clustering.
- Gene modules found by SciGeneX collectively reflect cell populations and their combinations are able to highlight specific cell populations, notably rare populations.
- SciGeneX showed high performance in selecting informative genes on artificial and experimental datasets and outperformed existing methods.
- The R package for SciGeneX is available for download at https://github.com/dputhier/scigenex.

## Methods

### The SciGeneX algorithm

The SciGeneX algorithm aims to identify and select modules of co-expressed genes. To do so, instead of examining cells in gene dimensions as in the conventional approach, SciGeneX analyzes genes in cell dimensions. In this space, genes whose expression is similar across cells are assumed to be close to each other. We illustrate genes in a 2D space to facilitate this visualization in Figure 1. From a more global point of view, sets of highly co-expressed genes corresponding to biological programs activated differentially between cells can generate dense regions of genes.

The SciGeneX algorithm aims to detect those groups of highly co-expressed biologically meaningful genes. It is divided in four main steps: (i) elimination of genes that do not co-express with other genes (ii) construction of a gene’s neighborhood graph (iii) identification of modules of co-expressed genes and (iv) filtering of modules of co-expressed genes.

In the initial step, the algorithm tries to identify the genes that are not near enough their neighbors to have a chance to co-express with them across the cells. To do so, the algorithm compares the distance of the nearest neighbors to the distribution of similar distances computed under the null hypothesis of random distribution of genes across the cell space. The genes having a distance too similar to the random distribution are removed from the downstream analysis. More precisely, the algorithm starts by computing a gene-gene distance matrix based, by default, on the Pearson correlation coefficient. This matrix reflects the relationships and similarities between different genes in terms of expression patterns. Alternative distance metrics are proposed in the package like Euclidean distances, Spearman correlation coefficient and cosine distances. Using this matrix, the algorithm computes the distance to the K^th^ nearest neighbors (DKNN) for each gene. This step quantifies the proximity of a gene to its closest neighbors in the gene expression space. The distribution of these distances for all genes is most often a long-tail distribution (Figure 1), suggesting the presence of co-expressed genes within the dataset. Next, to determine a threshold on DKNN above which genes will not be considered as co-expressing with other genes, SciGeneX computes the distribution of the DKNN distances under the hypothesis of no co-regulated genes. The algorithm uses random distance matrix permutations without replacement to compute null-hypothesis distance distribution and compares the observed distances to the simulated ones. This comparison aims to estimate the False Discovery Rate (FDR), which helps control the rate of false positives in the subsequent identification of co-expressed genes. From the simulated distances, SciGeneX determines a DKNN threshold below which distances are considered indicative of the presence of co-regulated genes. This threshold is determined based on the confidence level or the acceptable proportion of false positives. Genes with observed DKNN distances above this threshold are removed from the downstream analysis.

In the second step, the algorithm constructs a neighborhood graph where nodes represent genes. Edges in the graph can be defined using two methods. The first one, called *closest_neighborhood*, establishes an edge between genes A and B if B is part of the nearest neighborhood of size S for gene A (with S < K). The second one, called *reciprocal_neighborhood*, inspect the neighborhood of size K of all selected genes and put an edge between two genes A and B if they are reciprocally in each other’s neighborhood. For this last method, we recommend using a higher K value as more edges tend to be pruned using the reciprocity filter.

Finally, the algorithm searches to identify the densely connected regions of the neighborhood graph that may correspond to the highly-co-expressed genes across cells. To do so, the MCL algorithm is applied to the graph to create a partition of the genes and define gene modules. This algorithm simulates a random walk through the graph using expansion and inflation operators. This algorithm has successfully been used in various data analysis contexts including protein-protein interaction graph partitioning [47] or orthologous genes analysis [48], [49]. It is worth noting that MCL tends to produce several singleton clusters [50], thus, a filtering step, typically on module size or standard deviation of gene expression within a module, is subsequently applied.

The complete SciGeneX algorithm produces a list of gene modules, each module containing a set of genes that are highly co-expressed in cells and may correspond to biologically relevant processes that allow groups of cells to be distinguished by type or state.

### Implementation in R and analysis features

The SciGeneX algorithm has been implemented in a R package with a very easy-to-use API. The final co-expression gene modules are stored within an S4 object of the *ClusterSet* class, which has been implemented to offer functionalities for subsequent analyses. Users can utilize various features, such as (i) Filtering Options: filter based on criteria like gene number, gene names, or standard deviation, (ii) Effortless module selection: easily select gene modules using the indexing function (iii) Visualization capabilities: visualize results through interactive and non-interactive heatmaps, where cells are arranged using a hierarchical clustering algorithm (iv) Gene Ontology (GO) Enrichment Analysis: generate matrices for Gene Ontology (GO) enrichment analysis across co-expression modules (v) Reference Cell Markers: create matrices of reference cell markers that correspond to gene modules (vi) Global Expression Visualization: visualize the global expression of a gene module on the cell UMAP (vii) Export Options: export results to Excel spreadsheet or an HTML report for convenient data sharing. All these functionalities provide users with a comprehensive set of tools for in-depth exploration and interpretation of their data, enhancing the analytical capabilities provided by SciGeneX.

### Artificial datasets

We used the SPARSim R package to generate a comprehensive set of 100 scRNA-seq artificial datasets of 1755 cells distributed across 5 clusters with 7204 genes. Input parameters, including gene expression level intensities, gene expression level variabilities and sample library sizes, were estimated from the NK populations (n=155) of the PBMC3k dataset using the *SPARSim_estimate_parameter_from_data()* function. Gene expression level intensities were defined as the mean of the normalized counts across samples by the SPARSim methodology while, gene expression level variabilities as the variance of normalized counts across samples. The library size was established as the UMI count per cell. Within each dataset, we simulated 4 sets of DEGs, encompassing 500, 300, 200 and 50 genes, with a fold-change uniformly distributed between 4 and a maximum ranging from 10 to 100. The simulated parameters of DEGs (intensities, variabilities and library size) were estimated using the *SPARSim_create_DE_genes_parameter()* function. Thus, a total of 10 datasets for each 10 fold-change increment were generated using the *SPARSim_simulation()* function.

### Definition of true informative genes in Tabula Muris datasets

We downloaded the 12 droplet Seurat objects and their associated metadata from the Tabula Muris consortium website (https://tabula-muris.ds.czbiohub.org/). Each Seurat object contains single-cell transcriptomic data and represents a range of tissues including bladder, heart and aorta, kidney, limb muscle, liver, lung, mammary gland, marrow, spleen, thymus, tongue and trachea. We filtered out genes expressed in less than 3 cells for the subsequent analysis. Given the pre-defined cell populations, we identify the DEGs with the four distinct methods (*wilcoxon*, *t-test*, *bimod* and *MAST*) from the *FindAllMakers()* function (Seurat version 4.3.0) using default parameter except for only.pos = TRUE and min.pct = 0.25. Each method provided us with a set of true informative genes, enabling us to evaluate performance without relying on a single test. Genes were then selected based on a minimal average log2 fold-change of 0.5 and a maximum adjusted p-value of 0.05. These selected genes were considered the true informative genes for subsequent performance analysis.

### Evaluation of prediction performance

#### Artificial datasets

To predict the true informative genes in the artificial datasets, we applied both SciGeneX methods (*closest_neighborhood* and *reciprocal_neighborhood*) and two classically used methods from the Seurat package (VST and DISP). We pre-processed to datasets to only retain genes expressed in at least four cells. To select genes using the SciGeneX methods, we employed the *select_genes()* function with the following parameters: distance_method=“pearson,” noise_level=0.05, and a false discovery rate (FDR) of 1e-4. Notably, for the *closest_neighborhood* method, we used K=50, while we employed K=100 for the *reciprocal_neighborhood* method. Genes were subsequently clustered into co-expression modules using the *gene_clustering()* function. Graphs were partitioned using an inflation of 1.2. Finally, co-expression modules with fewer than 10 genes were filtered out using the *filter_cluster_size()* function. Modules containing less than 5 cells, expressing less than 20% of the module genes, were further excluded using the *filter_nb_supporting_cells()* function. Coupled with this, a selection of 2000 genes was done with the DISP and VST methods using the *FindVariableFeatures()* function from the Seurat package (version 4.3.0) with default settings. For sctransform method, genes were selected using the *SCTransform()* function from Seurat package (version 4.3.0). We used the *M3DropFeatureSelection()* function to predict informative genes with M3Drop (version 1.20.0), the *haystack()* and *show_result_haystack()* functions for singleCellHaystack (verison 1.0.2) and the *BigSur()* function for BigSur gene prediction (version 1.0.1). Subsequently, we computed the AUROC and F1-score for each method using the ROCR package (version 1.0-11). DKNN, variance, and dispersion values served as scoring metrics for gene ranking in AUROC calculations.

#### Tabula Muris datasets

The same methods were applied with slight parameter variations for the Tabula Muris datasets. In the same way as the artificial datasets, genes expressed in at least four cells were retained for the analysis. We selected genes with SciGeneX using the *select_genes()* function with parameters: distance_method=“pearson,” noise_level=0.05, and an FDR of 1e-8. For the *closest_neighborhood* method, we employed 30 neighbors, while for the *reciprocal_neighborhood* method, we used 100 neighbors. Graphs were generated with *gene_clustering()* function by establishing edges for the five closest neighbors (S) of each gene using the *closest_neighborhood* method. Conversely, the *reciprocal_neighborhood* method assigns edges a value of 0 for genes that do not fall within the neighborhood of their own neighbors. Subsequently, the graph partitioning was performed using an inflation parameter of 1.2. Co-expression modules with fewer than 10 genes were then filtered out using the *filter_cluster_size()* method. For these datasets, modules containing fewer than 5 cells expressing less than 35% were excluded using the *filter_nb_supporting_cells()* method. Finally, the methods used for comparison (DISP, VST, sctransform, M3Drop, singleCellHaystack and BigSur) were applied with the same functions and versions as in the artificial datasets section. Subsequently, we calculated the AUROC and F1-score for each method in the Tabula Muris datasets using the ROCR package. DKNN, variance, and dispersion metrics were used for gene ranking in the AUROC computations.

### PBMC3k dataset

#### Data preprocessing

The preprocessing step was performed with the Seurat R package (version 4.3.0). The *CreateSeuratObject()* method was employed to remove genes expressed in less than three cells as well as cells expressing less than 200 genes. An additional filter was applied to filter out cells expressing more than 5% of mitochondrial genes computed with the *PercentageFeatureSet()* method. Counts were then normalized using the *NormalizeData()* method with a scale factor of 10,000.

#### SciGeneX analysis

Genes were selected using the *select_genes()* method with the normalized count matrix as input. The following parameters were used: distance_method=“pearson,” noise_level=0.05, and an FDR of 1e-8. The subsequent genes were clustered into co-expression modules using the *gene_clustering()* function with the *closest_neighborhood* method. The graph was constructed with 3 nearest neighbors (S) for each gene and an inflation set to 1.8. The co-expression modules were then filtered to keep the ones containing at least 6 genes using the *filter_cluster_size()* method and the ones supported by at least 2 cells expressing more than 60% of genes in the co-expression modules using the *filter_nb_supporting_cells()* method. The top 20 highly co-expressed genes were selected using *top_genes()* method and heatmap of their expression were generated using the *plot_heatmap()* method in which cells were ordered by hierarchical clustering algorithm with ward.D as link.

#### GO enrichment analysis

GO enrichment analyses based on biological process terms were performed for each co-expression module using the *enrich_go()* method from the SciGeneX R package. Dotplot visualizations were generated using *viz_enrich()* and *plot_clust_enrcihments()* methods.

#### AUCell analysis

The AUCell R package (version 1.16.0) was used to highlight the fractions of enriched cells for each co-expression module. The ranking of cells was performed with the *AUCell_buildRankings()* method and AUC for each co-expression module in each cell was computed with the *AUCell_calcAUC()* method. Default parameters were used for both methods.

#### Kmeans clustering

Rare cell populations in Figure 4 were identified using the k-means clustering algorithm implemented in the amap R package (version 0.8-19). For co-expression module M08 and M11, euclidean distances were computed while for M15, manhattan distance was used.

#### ImmuNexUT analysis

We used the immune cell gene expression atlas, ImmuNexUT, to retrieve the immune cell types expressing specific co-expression modules in the PBMC3k dataset. This data contains 27 immune cell types (e.g plasmablasts and plasmacytoid dendritic cells) with their respective CPM values for x genes. CPM values were standardized using Z-score standardization and displayed for each gene in co-expression module M11 and M15 for every cell population present in ImmuNexUT database.

#### Cell cycle regression

As the co-expression module M08 was identified to be related to cell proliferation process, we used those genes to mitigate the effects of cell cycle heterogeneity in the PBMC3k dataset. Cell cycle regression was performed with the Seurat R package (version 4.3.0). A cell cycle score was first computed for each cell using the *CellCycleScoring()* method and was regressed out with the *ScaleData()* method. PCA was then performed using the informative genes identified by SciGeneX and the first 50 Principal Components were considered the UMAP dimension reduction representation.

### Human T cell trajectory dataset

The processed Human T cell trajectory dataset was downloaded on https://developmental.cellatlas.io website. Genes expressed in more than 100 cells were retained for the analysis. We used the normalized count provided to first identify co-expressed genes using *select_genes()* with parameters: distance_method=”pearson”, noise_level=0.25 and FDR=5e-4. The identified co-expressed genes are then clustered into co-expression gene modules using the *gene_clustering()* function with the *reciprocal_neighborhood* method and an inflation of 2. The subsequent gene modules were filtered to retain the ones containing more than 7 genes and with a minimal standard deviation of 0.3. Finally, we used the AUCell R package (version 1.16.0) to highlight the fractions of enriched cells for each co-expression module. Cells were ranked using the *AUCell_buildRankings()* function and AUC for each co-expression module in every cell was determined using the *AUCell_calcAUC()* function. We used the default parameters for both methods.

### Human thymic tissue section dataset

We used a public Visium spatial transcriptomics dataset of a human thymic section (from GEO Serie GSE207205) [41]. The preprocessing steps were performed using the Seurat package (version 4.3.0). The count matrix was log normalized using the *NormalizeData()* function and genes were scaled and centered using the *ScaleData()* function. A set of 2000 highly variable genes were selected with the *FindVariableFeatures()* function using the VST selection method and used for the principal component analysis. Spots were clustered based on the first 20 principal components using the *FindNeighbors()* and *FindClusters()*.

#### SciGeneX analysis

Genes expressed in at least 5 spots were kept for the analysis. Genes selection was performed on the normalized count matrix with the *select_genes()* function used with the following parameters: distance_method=“pearson” and row_sum=5. The *gene_clustering()* function was then called with the *closest_neighborhood* method and parameter set to: distance_method=”pearson” and inflation=2.2. Finally, those modules were selected to contain at least 7 genes and with a minimal standard deviation of 0.12. To quantify the expression level of each gene module, we computed module scores for each spot using the *AddModuleScore()* function from the Seurat package.

## Supporting information

Supp_file_Gene_Modules_Thymus_Single_Cell.xls

Supp_file_Gene_Modules_Thymus_Spatial_Transcriptomics.xls

## Acknowledgments

We thank Sébastien Jaeger, Romain Fenouil and Bertrand Escalière from the Computational, Biology, Biostatistics & Modeling (CB2M) platform and Delphine Potier (CRCM - Cancer Research Center of Marseille) for helpful discussions and advice, Sébastien Nin for his help to implement the package tests and Yann Kerdiles, Carole Siret and Isabella Bicalho Frazeto, (CIML - Centre d’Immunologie de Marseille Luminy) as well as Mathieu Adjemout (TAGC - Theories and Approaches of Genomic Complexity) for critical feedback and discussions.

## Data availability

The 12 10X genomics droplet datasets were downloaded as Seurat objects from the Tabula Muris web site (https://tabula-muris.ds.czbiohub.org). Annotations from the original studies were available in the metadata slot of the Seurat objects. The count matrix of the scRNA-seq PBMC3k dataset was obtained from 10x Genomics web site and can be downloaded from https://cf.10xgenomics.com/samples/cell/pbmc3k/pbmc3k_filtered_gene_bc_matrices.tar.gz. The scRNA-seq of thymic development dataset is available on ArrayExpress using accession number E-MTAB-8581 and was downloaded through the Human Cell Atlas Development portal (https://developmental.cellatlas.io/). The spatial transcriptomics dataset of human thymus was obtained from the Gene Expression Omnibus under accession number GSE207205. Only GSM6281326 was used.

## Code availability

Scripts and detailed instructions to reproduce the analysis for this manuscript are available on GitHub repository (link provided when accepted for review). A Docker image is also available on Zenodo (link provided when accepted for review) to easily reproduce the analysis.

## Authors contributions

DP and SvdP conceptualized, supervised the study and obtained funding. JB, JC and DP wrote the coding and analyzed the data. LS and SvdP provided intellectual input and conceptual advice. JB wrote the first draft. All authors read and approved the final manuscript.

