## Supplementary Data for "SciGeneX: Enhancing transcriptional analysis through gene module detection in single-cell and spatial transcriptomics data"

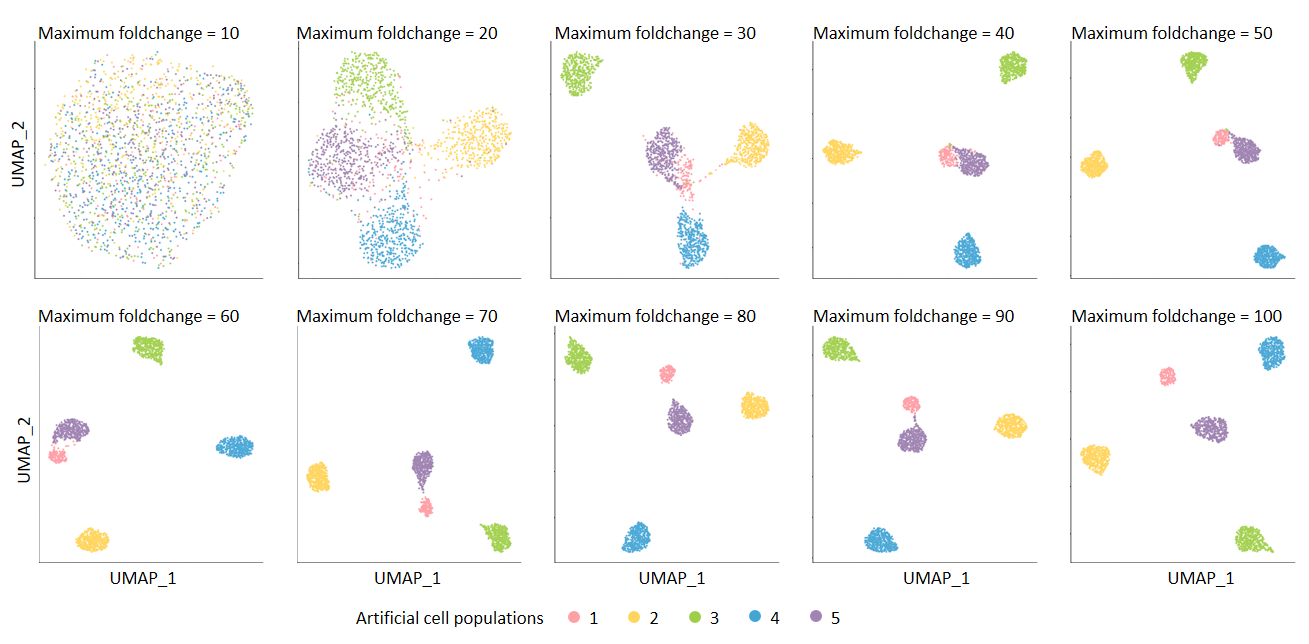

**Figure S1. UMAP representation of artificially generated scRNA-seq datasets.** UMAP representations of 10 artificially generated scRNA-seq datasets in which DEGs were simulated with maximum fold-changes ranging from 10 to 100 in increments of 10. Cell populations are indicated by distinct colors: purple, green, yellow, blue, and pink.

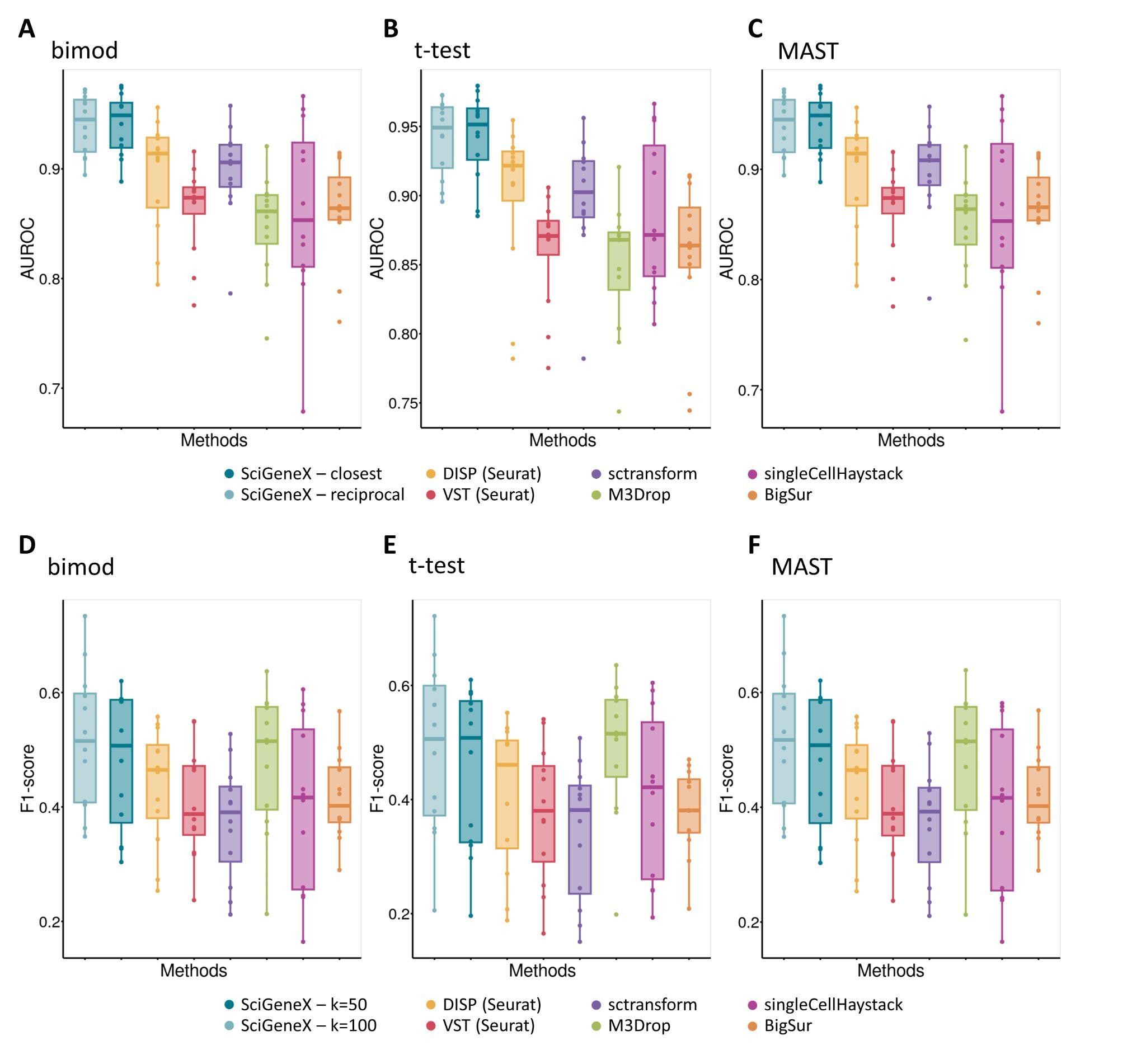

**Figure S2. Performance evaluation of different methods to find DEGs in Tabula Muris datasets.** Performances of SciGeneX and 6 existing methods were computed across a set of experimental datasets from the Tabula Muris consortium generated on 12 tissues. True DEGs have been defined using three methods, MAST (A, D), bimod (B, E) and t.test (C, F). AUROC (A, B, C) was used to evaluate the performances of SciGeneX neighborhood analysis for two k values, k=50 (dark blue) and k=100 (light blue) and F1-score (C, D, E) was used to evaluate both SciGeneX classifiers, *closest_method* (dark blue) and *reciprocal_method*. These results were compared to DISP (yellow), VST (red), sctransform (purple), M3Drop (green), singleCellHaystack (pink) and BigSur (orange).

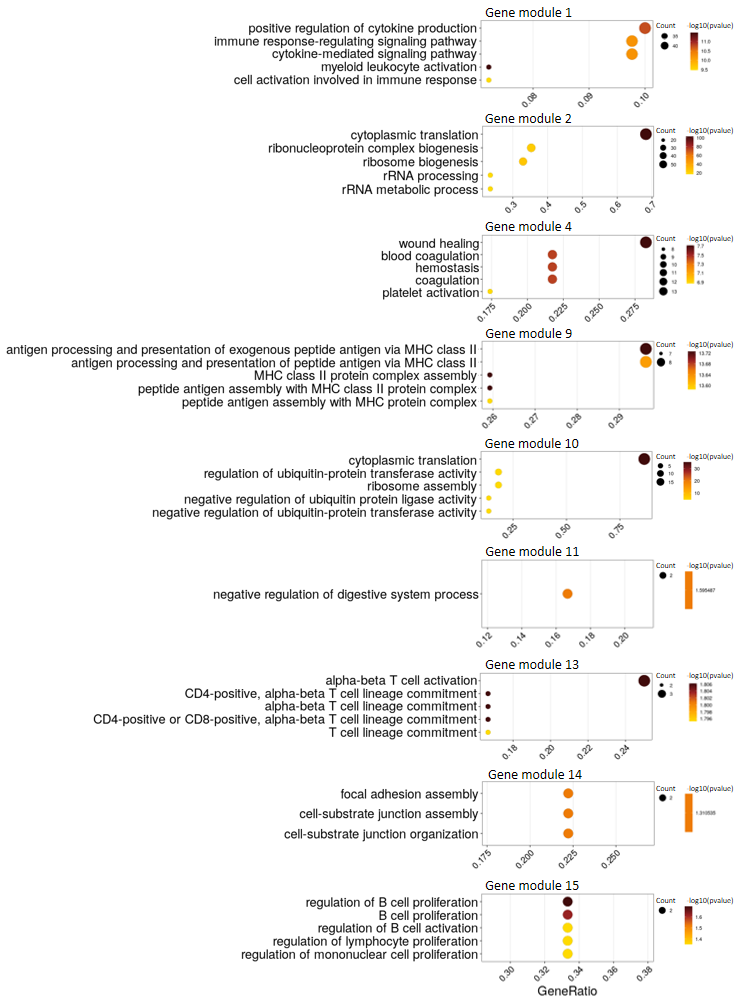

**Figure S3. Functional Enrichment Analysis of Co-Expressed Gene Modules Identified by SciGeneX.** Dotplot of functional enrichment analysis based on Gene Ontology (Biological Process) of co-expressed gene modules identified with SciGeneX in PBMC3k dataset. The y-axis represents the Gene Ontology terms, while the x-axis displays the gene ratio. Color represents the significance of their functional annotations.

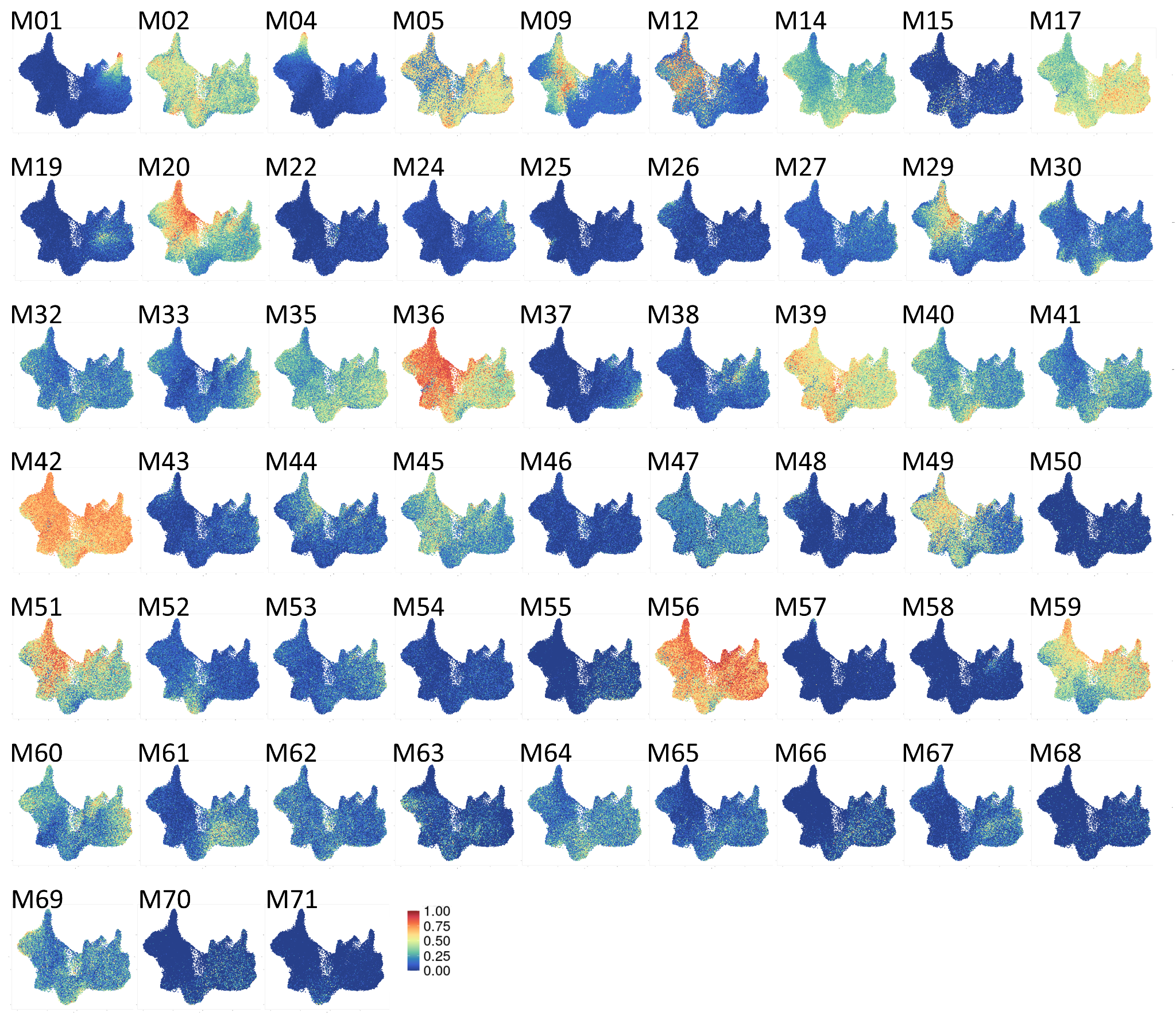

**Figure S4. UMAP visualization of AUCell scores for each gene module generated by SciGeneX in T cells trajectory dataset.** UMAP plots representing developing T cells from the T cell trajectory dataset. Cells are colored by the AUCell scores for each gene module generated by the SciGeneX algorithm.

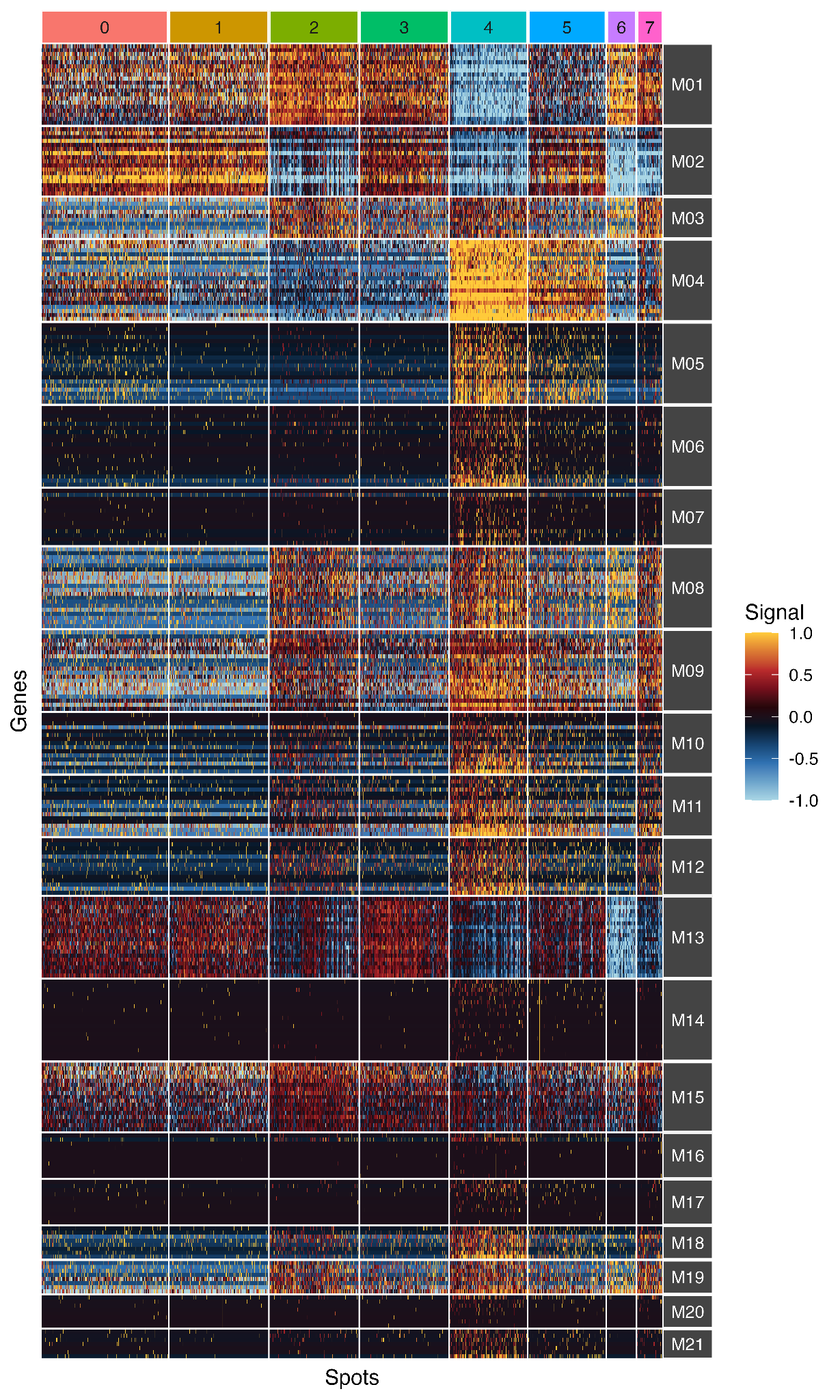

**Figure S5. Heatmap representation of co-expressed gene modules identified by SciGeneX in spatial transcriptomics of a human thymus section.** Heatmap displaying normalized expression levels of the top 20 genes within each co-expression module generated by the SciGeneX algorithm. The labels of the cell populations are shown on the top of the heatmap.

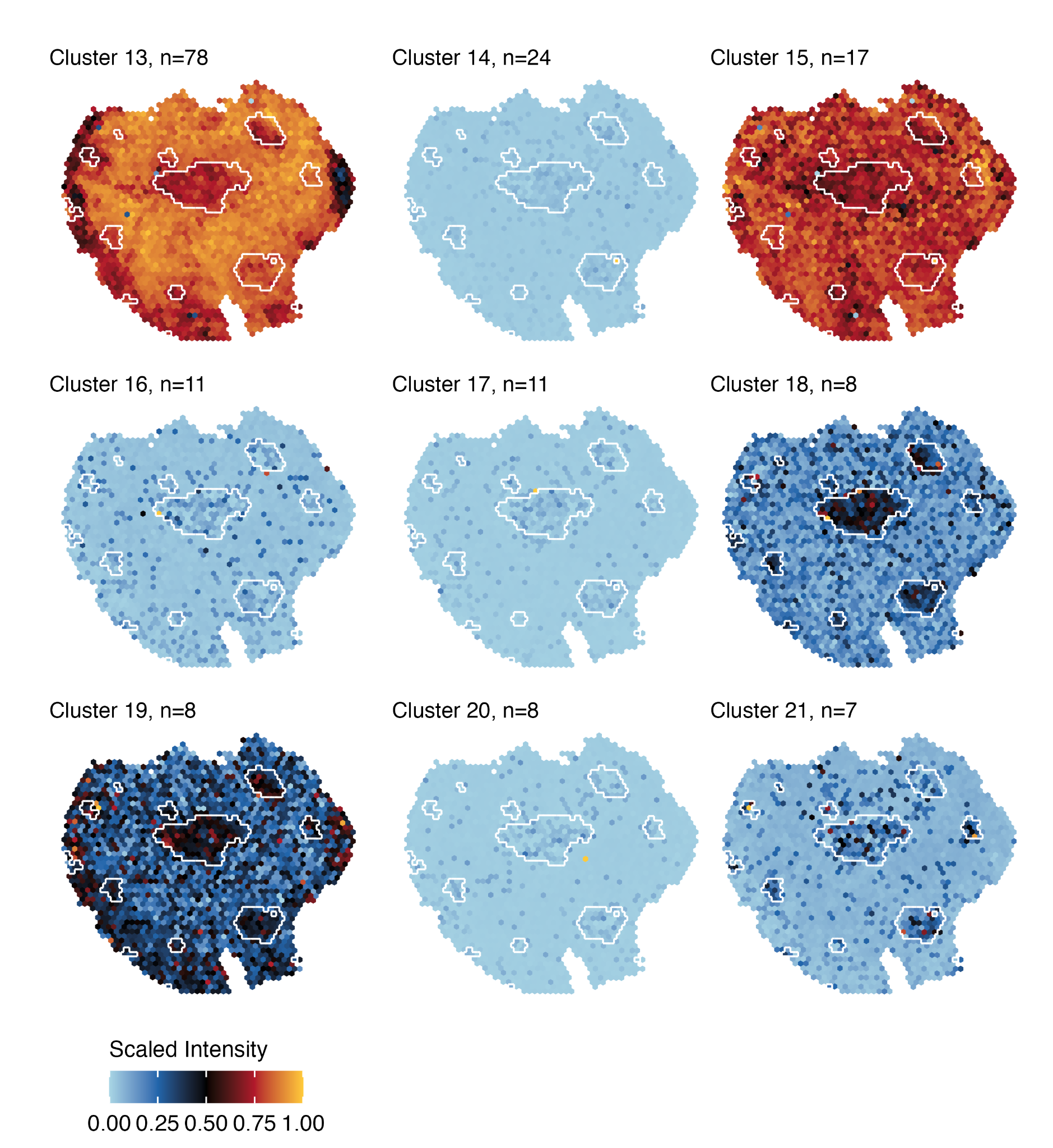

**Figure S6. Scaled expression intensity profiles of additional co-expressed gene modules on spatial transcriptomics dataset of a human thymus section.** Scaled expression intensity profiles of the co-expression modules not shown in Figure 6. Each spot on the spatial transcriptomics map represents the scaled intensity of co-expressed gene modules within the human thymus section.

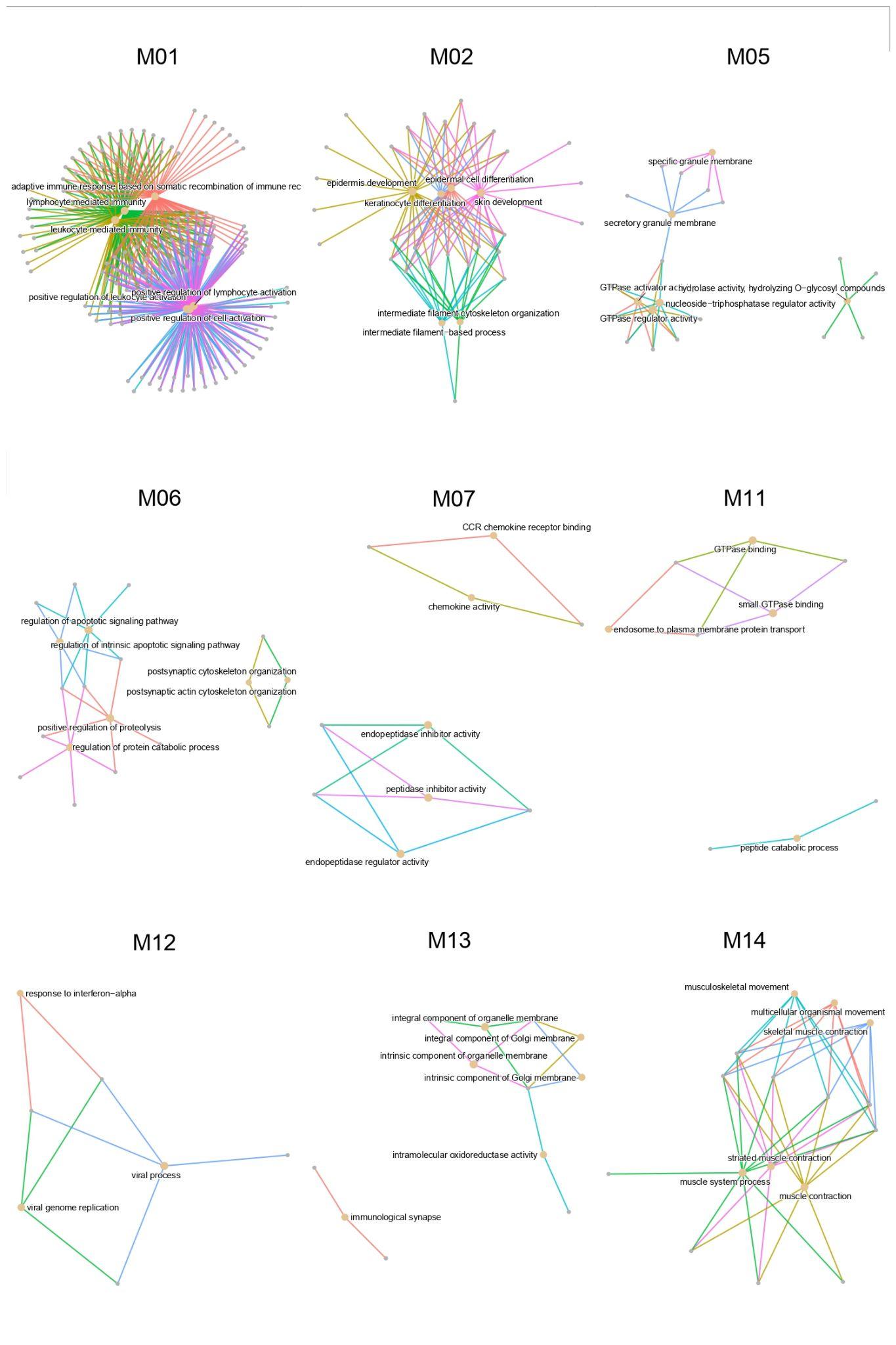

**Figure S7. Network representation of functional enrichment analysis of co-expressed gene modules obtained on spatial transcriptomics dataset of a human thymus section.** Network representation of the results obtained from the functional enrichment analysis based on Gene Ontology (Biological Process) of co-expressed gene modules identified within the spatial transcriptomics dataset of a human thymus section. The network visually represents the enriched biological processes associated with these gene modules.
